# Mosquito species and age influence thermal performance of traits relevant to malaria transmission

**DOI:** 10.1101/769604

**Authors:** KL Miazgowicz, EA Mordecai, SJ Ryan, RJ Hall, J Owen, T Adanlawo, K Balaji, CC Murdock

**Author notes:** Corresponding Author: Courtney C. Murdock.

## Abstract

Models predicting disease transmission are a vital tool in the control of mosquito populations and malaria reduction as they can target intervention efforts. We compared the performance of temperature-dependent transmission models when mosquito life history traits were allowed to change across the lifespan of *Anopheles stephensi*, the urban malaria mosquito, to models parameterized with commonly derived estimates of lifetime trait values. We conducted an experiment on adult female *An. stephensi* to generate daily per capita values for lifespan, egg production, and biting rate at six constant temperatures. Both temperature and age significantly affected trait values. Further, we found quantitative and qualitative differences between temperature-trait relationships estimated based on daily rates versus directly observed lifetime values. Incorporating these temperature-trait relationships into an expression governing transmission suitability, relative *R*_0_(*T*), model resulted in minor differences in the breadth of suitable temperatures for *Plasmodium falciparum* transmission between the two models constructed from only *An. stephensi* trait data, but a substantial increase in breadth compared to a previously published model consisting of trait data from multiple mosquito species. Overall this work highlights the importance of considering how mosquito trait values vary with mosquito age and mosquito species when generating temperature-based environmental suitability predictions of transmission.

## Introduction

Despite the progress of global malaria elimination programs in reducing the incidence of human malaria, particularly *Plasmodium falciparum*, malaria remains a leading cause of morbidity and mortality among infectious diseases (1). The occurrence of multi-class drug and insecticide resistance, in addition to alterations in mosquito behavior, challenge our ability to eradicate malaria and poses the possibility of resurgence (1–6). While numerous environmental factors affect the distribution and prevalence of mosquito-borne diseases, temperature is one of the most pervasive abiotic factors affecting both mosquito and parasite vital rates (7–23). However, even though the importance of these factors is increasingly recognized, gaps remain in the current mechanistic understanding of the relationship between malaria risk and key environmental variables. Improving our understanding of the link between temperature and malaria transmission will be crucial for predicting how transmission varies geographically, seasonally, and with climate and land use change (10, 24–30).

Recent research has begun to explicitly define the relationship between temperature and vector and pathogen traits relevant to transmission across a diversity of vector-borne disease systems (24, 26, 31–37). The net effect of these traits in determining temperature-dependent transmission potential can be described by the pathogen basic reproduction number (*R*_0_), defined as the number of secondary cases arising from a primary case given a fully susceptible population. Transmission models that define *R*_*0*_ can be used to generate predictions of disease risk, inform intervention strategies, and evaluate efficacy of various disease interventions (38–43). Although it is widely accepted that the life history traits of ectotherms exhibit unimodal responses to temperature, models often assume that the temperature-trait relationships for key mosquito and parasite life history traits are linear (37, 44–48). Further, recent research that included nonlinear, unimodal temperature-trait relationships into an *R*_*0*_ model for malaria transmission predicted a lower temperature optimum, minimum, and maximum than previous estimates based on linear relationships, which better matched field observations of entomological inoculation rate (31, 32). Key biological insights from previous models are that: 1) malaria transmission is constrained in hot summer months in equatorial and tropical regions, 2) regions of the world that are currently permissive for transmission may become less environmentally suitable with future climate warming, and 3) vector control may become more difficult as temperatures in northern latitudes become more permissive and suitable seasons extend (24, 32, 49).

Despite these advances, insights from previous mechanistic *R*_*0*_ models remain constrained by a lack of entomological and parasite data (31). Temperature-trait relationships for key parameters are often indirectly estimated from a limited number of studies, leading to high uncertainty around the predicted thermal limits in current malaria *R*_*0*_ models (31, 32). Additionally, the parameterization of *R*_*0*_ models with temperature-trait relationships aggregated from different mosquito and parasite species likely introduces error and uncertainty in *R*_*0*_ estimates given the degree of inter- and intra-species variation in life history (7, 31, 32).

Further, evidence from a diversity of invertebrate systems demonstrates that organisms experience age-related changes in life history traits (50–53). These changes reflect either senescence, a decline in general physiological function with age, or a shift in energy allocation to different life history tasks as an organism ages to maximize fitness. Limited studies suggest that age modifies mosquito life history, with some evidence that mosquitoes experience reproductive senescence (54), bite more frequently as they age (55), and exhibit age-dependent survivorship (52). Yet, incorporating the combined effect of temperature and age on mosquito life history traits has not been explicitly addressed in temperature-dependent *R*_*0*_ models for malaria. Studies often use data collected over a relatively limited portion of the mosquito lifespan to estimate lifetime traits such as biting rates, total egg production, and lifespan in models of mosquito population dynamics and disease transmission (23, 24, 26, 31, 32, 35, 36). For example, a recent study characterized the extrinsic incubation period of *P. falciparum* in a cohort of *An. stephensi* mosquitoes, but used the duration of the gonotrophic cycle to approximate the daily biting rate and thus force of infection (23). If key mosquito life history traits vary with age, and temperature influences age-related changes in these traits, then precisely when these traits are measured during the lifespan of the mosquito could impact the predicted relationships between these traits and temperature as well as the predicted environmental suitability for malaria transmission.

In this study, we conducted a cohort life table experiment on the urban Indian malaria vector (*An. stephensi*) at six different constant temperatures to address the following questions: 1) does the effect of temperature on transmission change when thermal responses of traits from a single mosquito and parasite species are used, rather than aggregated from multiple mosquito and parasite species? 2) how do *An. stephensi* life history traits vary across the full spectrum of biologically relevant temperatures? 3) do life history traits that drive human malaria transmission vary with mosquito age? and if so, 4) do age-dependent changes in life history affect temperature-trait responses and the overall temperature suitability for transmission?

## Materials & Methods

### Life history experimental design

*An. stephensi* mosquitoes were reared as described in **SI_Methods**. The life history experiment was initiated three days after adult emergence to permit mating. After they were presented with an initial blood meal for 15 minutes via a water-jacketed membrane feeder we randomly distributed 30 host-seeking females into individual cages (16 oz. paper cup with mesh top) to one of six constant temperature treatments (16°C, 20°C, 24°C, 28°C, 32°C, 36°C ± 0.5°C, 80% ± 5 RH, and 12L:12D photoperiod). Each individual adult cage contained an oviposition site: a small petri dish that secured to the bottom of the housing, containing cotton balls to retain liquid, overlaid with a filter paper for easy egg removal and counting. Individual mosquitoes were offered a blood meal for 15 min each day. An individual was scored as having taken a blood meal through visual verification of the abdomen immediately after the feeding period. Oviposition sites were rehydrated and checked for the presence of eggs daily. We followed populations of individual females in each temperature treatment until all mosquitoes had died or when less than 7% of the starting population remained. At least two biological replicates were performed at each temperature (n= 30 per temperature; total n= 390 individuals). Given the extended duration of these experiments (~60 days), multiple blood donors were used throughout each experimental replicate.

### Statistical analysis

All statistical analyses were performed using the program R (version 3.4.1). We used generalized linear mixed models (GLMM) R package <*lme4∷glmer()*> to estimate the effects of temperature, mosquito age, and their interaction on the proportion of females that imbibed blood on a given day (i.e., the number of females that took a blood meal on a given day out of the total number of females alive on that day for each temperature treatment) and the mean daily egg production (i.e., the number of eggs laid on a given day divided by the total number of females alive on that day in a given temperature treatment) (56). Temperature, age, and their interaction were included as fixed effects. Both temperature and age were included as continuous variables that were scaled and centered. Random factors initially included block, blood donor, and mosquito individual as categorical variables. We used minimum AIC values to compare and select our final models (**SI_Table 1, SI_Table 2**). Second, we used a Log-rank test with R package <*survival∷survdiff()*> on Kaplan-Meier estimates to determine if survivorship differed with temperature (57). Lastly, to determine if the daily survival rate changed across the lifespan of the mosquito, we fit a variety of survival distributions, which allow either for a constant (exponential) or variable daily mortality rate (log-normal, gamma, Gompertz, and Weibull) with R package <*flexsurv*> to the Kaplan-Meier estimates (58). AIC values were used to determine survival distribution fits. In addition, the best fitting distributions (minimum AIC) were determined at each temperature treatment separately to confirm the best-fitting survival distribution did not vary with temperature treatment (**SI_Table 3)**.

### Temperature-dependent transmission potential (R_0_)

We used a temperature-dependent *R*_*0*_ framework parameterized from the *An. stephensi-P. falciparum* system to 1) compare predicted environmental suitability for malaria transmission to a previous *R*_*0*_(*T*) model that aimed to describe the *An. gambiae* – *P. falciparum* system but consisted of data aggregated from several different mosquito and parasite species (31); and 2) to evaluate the effect of age-related changes in *An. stephensi* life history on the predicted environmental suitability of *P. falciparum*. To do this, we used a common expression for *R*_*0*_ derived from the Ross-MacDonald model (41, 59), which was initially expanded on in Parham & Michael (2010) to incorporate the effect of temperature on mosquito life history and thereby mosquito population size, and later modified in Mordecai et al. (2013) to approximate individual lifetime reproductive values using daily fecundity output and adult daily mortality rates. (**Equation 1**, **SI_Table 4**) (24, 26, 31–37, 59):

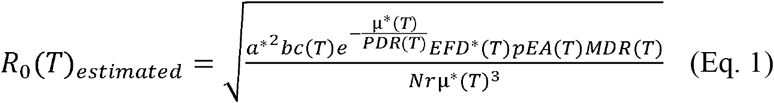

*R*_*0*_ is the expected number of new cases generated by a single infectious person or mosquito introduced into a fully susceptible population throughout the period within which the person or mosquito is infectious. *R*_*0*_ components include: egg-to-adult survival probability (*pEA*), mosquito development rate (*MDR*), fecundity (*EFD;* eggs laid per female per day), biting rate (*a*), adult mosquito mortality rate (*μ*), parasite development rate (*PDR*), vector competence (*bc*; the proportion of parasite-exposed mosquitoes that become infectious), the density of humans (*N*), and the human recovery rate (*r*), with (*T*) indicating parameters that are dependent on environmental temperature (degrees Celsius). The host recovery rate (*r*) and host density (*N*) are assumed to be temperature independent. We label the *R*_*0*_*(T)* formulation in **Eq. 1** as ‘estimated’ as lifetime traits are commonly parameterized with indirect estimates based on daily rates (24, 26, 31–36).

To construct the Multi-species estimated model (*R*_*0*_*(T)*Multi-species estimated) we used the thermal relationships defined in (31) and using the formulation in **Eq. 1**. To compare the Multi-species estimated model to the *R*_*0*_*(T)* model parameterized with our *An. stephesnsi* data (*R_0_(T) An. stephensi* estimated) and using the formulation in **Eq. 1**, we generated trait estimates (denoted by *) according to methods described in (13, 22, 31, 32) for biting rate (a*), lifespan (lf* as 1/ *μ* *), and lifetime egg production (B* as EFD*/*μ* *). Briefly, the inverse of the duration of the first gonotrophic cycle for each individual was used to estimate biting rate (*a*\*). Exponential curves were fit to the tail of mosquito survivorship distributions characterized at different constant temperatures to estimate the daily mortality rate (*μ*\*) of mosquitoes at each temperature treatment. Eggs laid per female per day (*EFD*\*) at each temperature was estimated by dividing the number of eggs laid for each female in her first gonotrophic cycle by the number of days in that gonotrophic cycle. Additionally, to estimate *An. stephensi* mosquito development rate (*MDR*) and probability of egg to adult survival (*pEA*), as well as *P. falciparum* development rate (*PDR*) and vector competence (*bc*), we used data from (11) and (23). Finally, to incorporate the temperature-dependence (*T*) of each of the traits outlined above and below, we fit symmetric and asymmetric non-linear responses using Bayesian inference as described in Johnson et al. 2015 and **SI_Methods** (31).

To determine if *R*_*0*_*(T)* varies when directly observed lifetime trait values for biting rate (*a*), lifespan (*lf*), and lifetime egg production (*B*) are incorporated instead of indirect estimates of the lifetime values for these traits for *An. stephensi*, we generated the following *R*_*0*_*(T)* formulation (**Equation 2**, **SI_Table 4**).

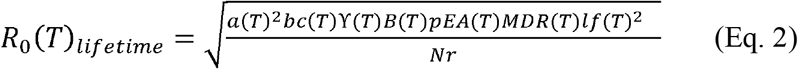

Mosquito lifespan (*lf*) was defined as the total number of days a mosquito survives after being placed within her respective temperature treatment. Individual lifetime biting rate (*a*) was defined as the total number of blood meals a female takes during her lifespan (*lf*) divided by her lifespan (*lf*). Lifetime egg production (*B*) is defined as the total number of eggs laid by a female during her lifespan (*lf*). The directly observed biting rate (*a*), lifespan (*lf*), and lifetime egg production (*B*) were substituted for the indirectly estimated biting rate (*a*\*), lifespan (*lf*\* = 1/ *μ*\*), and lifetime egg production (*B*\* = *EFD*\*/*μ*\*) in **Eq. 1**. The proportion of mosquitoes surviving the latency period, denoted as ◻ in **Eq. 2,** is substituted for exp[-*μ/PDR*] in **Eq. 1**. To estimate ◻, we first fit a Gompertz distribution to survivorship data from each temperature treatment and experimental replicate. We then took the proportion of mosquitoes alive upon completion of the predicted extrinsic incubation period (*PDR_50_*(*T*)^−1^) of *P. falciparum* at each temperature. The amount of days to reach 50% of maximum infectiousness in a mosquito population is represented by *PDR_50_*(*T*)^−1^ (23). This formulation allows us to account for age-dependent mortality in the proportion of mosquitoes surviving the latency period (◻). We then compared the relationship lifespan, biting rate, and lifetime egg production have with temperature for *An. stephensi* when these traits are directly observed (*lf*, *a*, *B*) vs. estimated (*lf*, a*, B\**) from the data generated in this study, as well as if any observed differences translate to differences in the predicted environmental suitability for malaria transmission (*R*_*0*_).

As done previously (24, 26, 31–36)), we use relative values of *R*_*0*_, as opposed to absolute values, to estimate relative temperature suitability for malaria transmission, because absolute values of *R*_*0*_ depend on a number of factors that vary by location and time, including humidity, breeding habitat availability, vector control, and vector – human contact rates. By rescaling *R*_*0*_*(T)* in all models to a range between 0 and 1, we can easily compare the thermal optimum and limits for relative *R*_*0*_ across models. However, when adopting this relative approach, the stable transmission threshold of *R*_*0*_ >1 is no longer meaningful. Therefore, a conservative suitability threshold of relative *R_0_(T) >0* is implemented where temperatures outside of this range are deemed unsuitable for transmission to occur because one or more of the component traits in *R*_*0*_*(T)* is equal to zero. Finally, sensitivity and uncertainty analyses were performed for our *An. stephensi* estimated and lifetime models as described in (31) and **SI_Methods**.

### Mapping seasonal transmission range

We generated maps depicting the number of months an area is predicted to be environmentally suitable for transmission of human malaria (*P. falciparum*) to illustrate the potential impact differences in the thermal breadth among our relative *R*_*0*_*(T)* models have across a relevant landscape. We were primarily interested in comparing the amount of area predicted to be endemically (year-round) suitable for *P. falciparum* transmission between our two *An. stephensi* relative *R*_*0*_*(T)* models incorporating either estimated or observed lifetime trait values across the current distribution of *An. stephensi*. However, we also draw comparisons between our *An. stephensi* models (*An. stephensi* estimated and lifetime) and a previously derived relative *R*_*0*_*(T)* model (Multi-species estimated), which aggregates trait data from multiple mosquito and parasite species intended to describe *P. falciparum* transmission in the *An. gambiae* system. This latter comparison serves to illustrate the differences in temperature suitability among systems. Using the median model output for each of these models, we calculated *R*_*0*_*(T)* values at 0.2°C increments, at a 0.01 level accuracy of model output, rescaled the *R*_*0*_*(T)* values from 0-1, and plotted transmission suitability where *R*_*0*_*(T)* >0 as in Tesla et al. 2018 (35). Using the GADM global administrative boundaries data we estimated the land area with endemic (year-round) suitability within countries that span the current range for *An. stephensi*: India, Pakistan, Sri Lanka, Qatar, United Arab Emirates, and Oman (60).

## Results

### Temperature and age shape mosquito life history traits

A cohort life table study was used to evaluate the effect of temperature on *An. stephensi* life history traits as individuals age. Both temperature and mosquito age significantly affected the proportion of females that imbibed blood on a given day, mean daily egg production, and survivorship (**Figure 1**, **SI_Table 2**). However, the interaction between temperature and age did not significantly affect the proportion of females that imbibed blood on a given day or mean daily egg production (**SI_Table 2**). The proportion of females that imbibed blood on a given day was generally higher at warmer temperatures and declined as mosquitoes approached the end of their lifespan in all temperature treatments, with this age-associated decline being most pronounced at 36°C (**Figure 1a**). Across all temperature treatments, mean daily egg production increased over time to a peak value before declining (**Figure 1a).** Peak mean daily egg production varied with temperature: peak values were higher, occurred sooner, and persisted for shorter periods of time at warmer temperatures (28°C - 36°C) compared to cooler temperatures (16°C - 24°C) (**Figure 1b**). Temperature also significantly affected survivorship (**SI_Table 2**). Survival responded unimodally to temperature, with a peak at 20°C and a decline at higher temperatures (**Figure 1b**). Finally, at all temperatures, mosquito daily probability of survival was not constant with age: a Gompertz distribution, which allows for a variable daily mortality rate, best fit the survival data at each temperature (**Figure 1c**, **SI_Table 2**).

**Figure 1.**
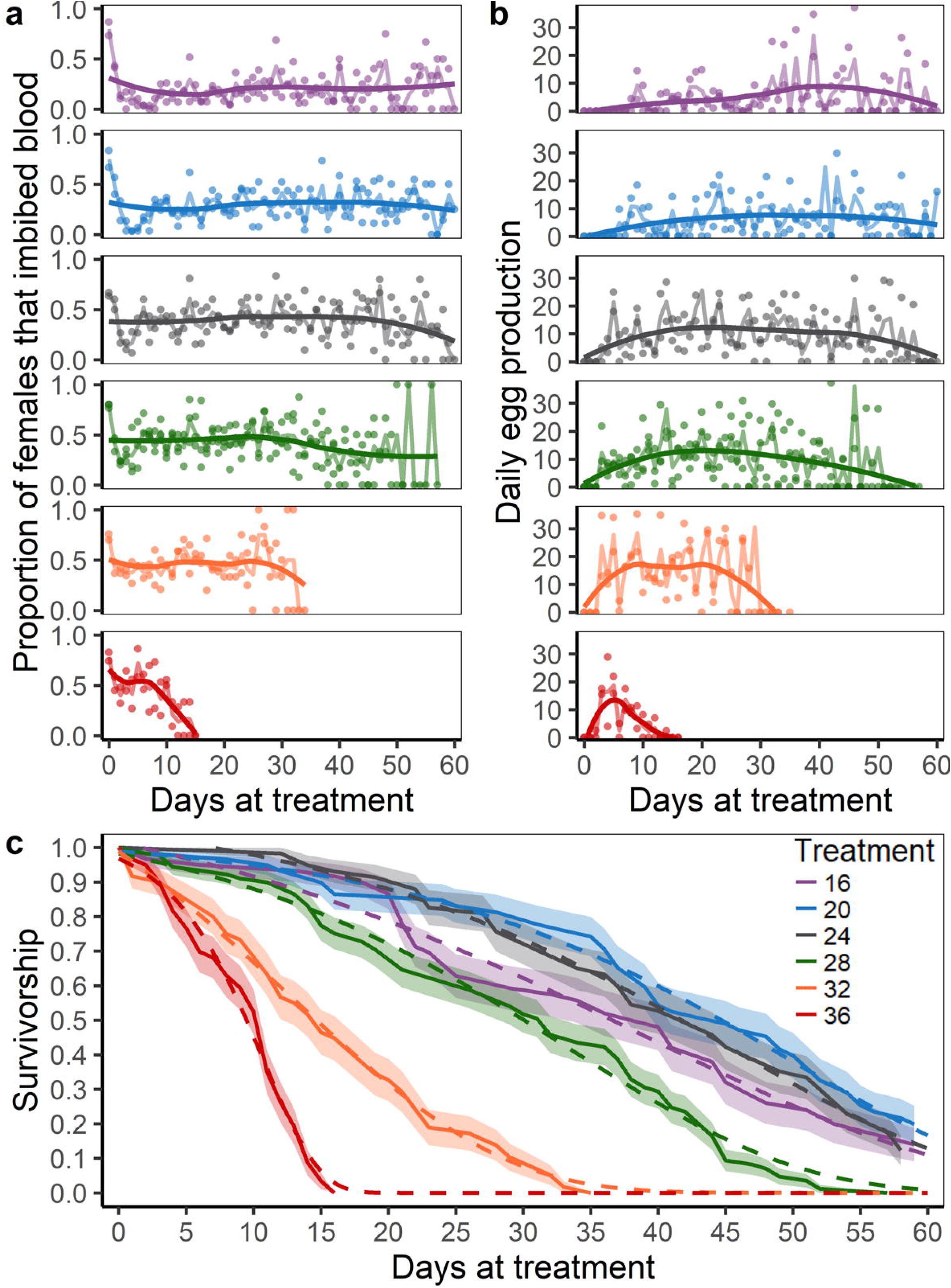
Temporal Effects. *Anopheles stephensi* life history daily trait values across six temperature treatments (16°C, 20°C, 24°C, 28°C, 32°C, and 36°C) for the (**a**) proportion of females that imbibed blood, (**a**) daily egg production, and (**c**) survivorship. In **a** and **b**, colored dots represent trait values for each replicate, with smoothed averages (loess; solid line) and the trend across daily mean values (faded solid line) shown. The proportion of females that imbibed blood, is the defined as the total number of females who took a blood meal on a given day over the total number of females alive on that day. Daily egg production is defined as the total number of eggs laid on a given day over the total number of female alive on that day. In **c**, Kaplan-Meier estimates (solid line, upper and lower 95% CI: shaded area) and the best-fitting survival distribution (Gompertz; dashed line) are shown.

### Using directly observed as opposed to estimated lifetime trait values alters temperature-trait relationships

Depending on the life history trait examined (biting rate (*a*), lifespan (*lf*), or lifetime egg production (*B*)), using observed vs. estimated lifetime values to fit temperature-trait relationships resulted in shifts in the predicted thermal minimum (*T*_*min*_), maximum (*T*_*max*_), and optimum (*T*_*opt*_) (**Figure 2a-c**, **SI_Table 5**). Temperature-trait relationships derived from estimated lifetime trait values resulted in an overall decrease in the absolute values for each lifetime trait (**Figure 2a-c**). While peak values of the temperature functions for observed lifetime biting rate (*a*) were approximately double (0.5 vs. 0.24) that of estimated lifetime biting rate (*a*\*), the temperature at which these peak values occurred (*T*_*opt*_) was 4.2°C lower for observed lifetime biting rate (**Figure 2a**, **SI_Table 5**). Further, the temperature-trait relationship for estimated biting rate (*a*\*) had a substantially warmer predicted thermal minimum (*T*_*min*_; +7.6°C) and maximum (*T*_*max*_; +2.4°C) than that for lifetime biting rate (*a*) resulting in a 5.2°C reduction in the breadth of temperatures (*T_breadth_*) permissive for biting (**Figure 2a**, **SI_Table 5**). Similarly, the value for observed lifespan (*lf*) at the predicted thermal optimum (*T*_*opt*_) was approximately twice that of estimated lifespan (*lf*\*; 38.3 days vs. 16 days) (**Figure 2b**, **SI_Table 5**). However, in contrast to biting rate, the predicted optimum and maximum temperatures were the same for observed lifespan (*lf*) and estimated lifespan (*lf*\*) with only a slight 0.2°C difference in the predicted thermal minimum (**Figure 2b, SI_Table 5**). The temperature-trait relationship for observed lifetime egg production (*B*) was predicted to have a 2.2°C decrease in the *T*_*opt*_ as compared to estimated lifetime egg production (*B*\*), with subtle shifts in the predicted thermal minimum and maximum (**Figure 2c**, **SI_Table 5**). Predicted peak values were higher for observed lifetime egg production (*B*; 317.1 eggs) than estimated lifetime egg production (*B*\*; 225 eggs) (**Figure 2c**).

**Figure 2.**
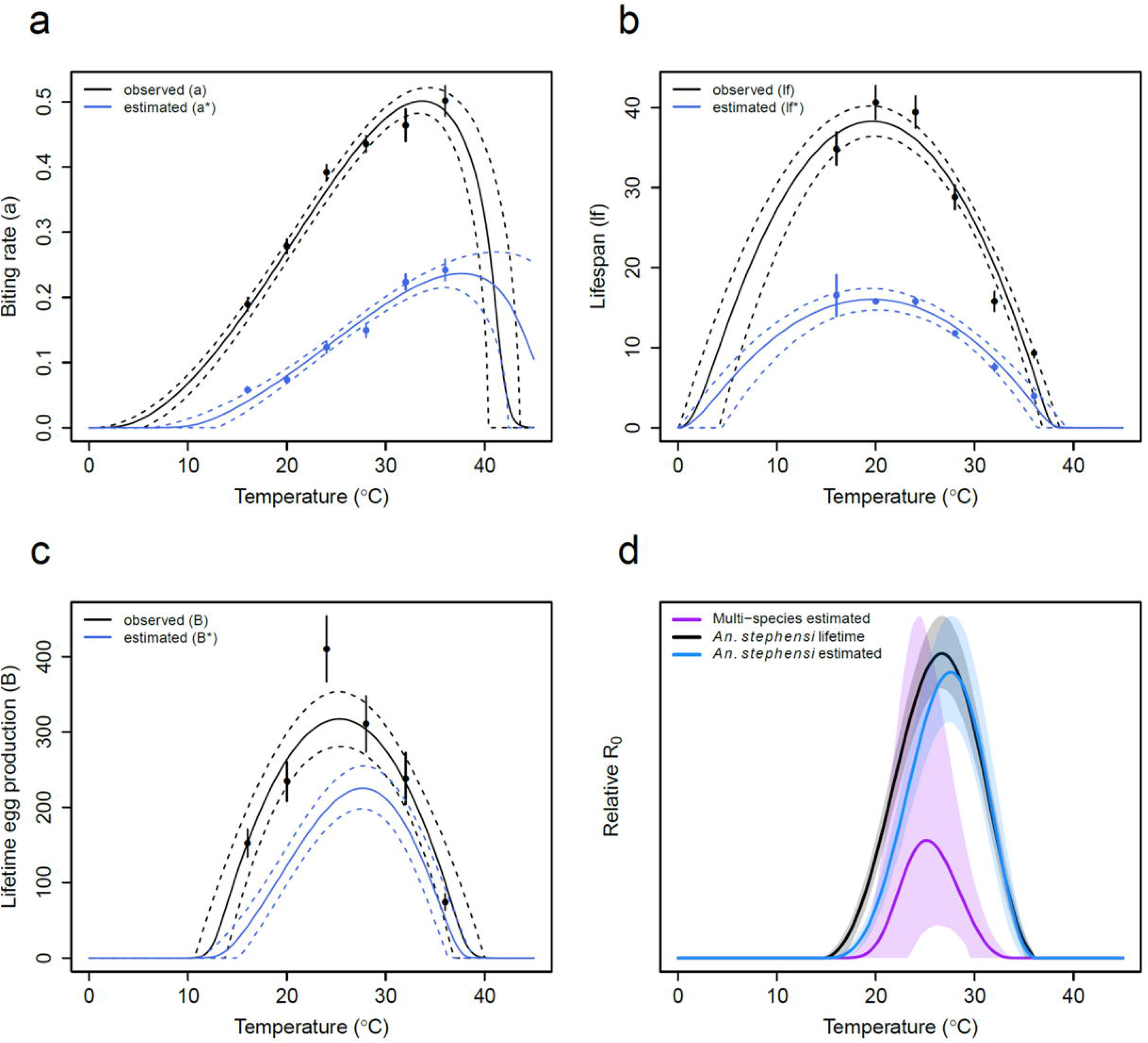
Direct Comparison of Lifetime vs. Estimated Traits. A direct comparison of temperature-trait relationships between observed lifetime trait values (black) and estimated lifetime values (blue) for (**a**) biting rate (*a*), (**b**) lifespan (*lf*), and (**c**) lifetime egg production (*B*). No data points (dots) are displayed for estimated lifetime values in **c**, as this trait is the product of the temperature-trait relationships for estimated daily egg production and estimated lifespan (*B*(T)* = *EFD*(T) x lf*(T)*). Comparison of the three relative *R*_*0*_*(T)* models (**d**). Each model is displayed as relative to the respective max *T*_*opt*_ upper 95% credible interval (CI) model value with mean model values (solid line) and 95% CI (faded area) across temperature shown.

Surprisingly, the changes in temperature-trait relationships that occurred when directly observed lifetime data are used instead of indirect estimates did not yield large changes in the predicted relationship between temperature and malaria transmission across the *An. stephensi* relative *R*_*0*_*(T)* models (**Figure 2d**, **SI_Table 6**). There was a slight decrease from 27.6°C (*An. stephensi* estimated) to 26.6°C (*An. stephensi* lifetime) in the predicted *T*_*opt*_, but no major differences noted in the predicted *T*_*min*_ and *T*_*max*_ across models. We did also find notable differences in the *T*_*min*_ and *T*_*max*_ of the estimated thermal relationship for the proportion of mosquitoes surviving the latency period (◻) between *An. stephensi* models, which also likely contributed to the minor shifts in relative *R*_*0*_*(T)* (**SI_Figure 4, SI_Table 7**). Finally, *R*_*0*_*(T)* was sensitive to lifespan (*lf*) and biting rate (*a*) in both *An. stephensi* models; however, the *An. stephensi* lifetime model exhibited less sensitivity to lifespan (*lf*) than the *An. stephensi* estimated model (**SI_Results**, *An. stephensi* lifetime; **SI_Figure 2** & *An. stephensi* estimated; **SI_Figure 3**).

**Figure 3.**
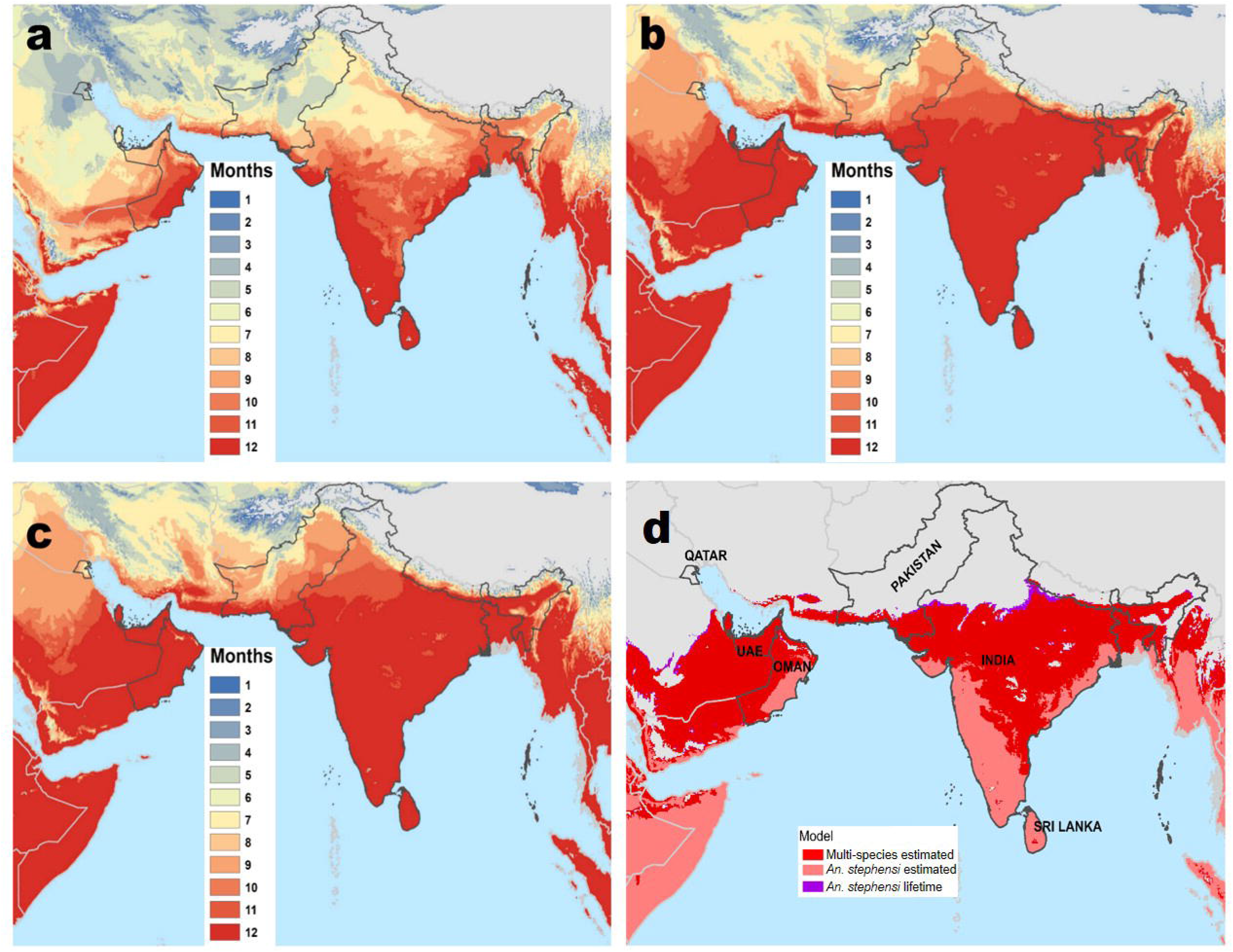
Mapping of relative *R*_*0*_*(T)*. Number of monthly pixels with *R*_*0*_*(T)*>0 for (**a**) Multi-species estimated, (**b**) *An. stephensi* estimated, and (**c**) *An. stephensi* lifetime. (**d**) Mapping overlay of all three models with endemic transmission (12-months suitability, deep red shading in previous panels) with *An. stephensi* lifetime (bottom layer, purple), *An. stephensi* estimated (middle layer, deep red), and Multi-species estimated (top layer, pink). Thus, the pink region corresponds approximately to the areas where all three models predict endemic suitability, the deep red reflects the additional area predicted to have endemic suitability by the *An. stephensi* models, while the purple represents the additional area predicted for endemic suitability by the *An. stephensi* lifetime model only.

### The relationship between temperature and relative R_0_ is disease system specific

Integrating temperature-trait relationships from the *An. stephensi – P. falciparum* system resulted in a qualitatively different temperature-relative *R*_*0*_ relationship to a previously defined multi-species model (31) (**Figure 2d**, **SI_Table 4, SI_Table 6**). The *An. stephensi* relative *R*_*0*_(*T*) model parameterized with equivalent methods (*An. stephensi* estimated) displayed an increase in the breadth of suitable temperatures over which *R*_*0*_ > 0 and a decrease in the credible intervals around the thermal minimum (*T*_*min*_), maximum (*T*_*max*_), and optimum (*T*_*opt*_) compared to a previously published model (Multi-species estimated) which was used to describe *P. falciparum* transmission via *An. gambiae* (**Figure 2d, SI_Table 6**). This increase in the range of temperatures that are suitable for malaria transmission results from an increase in *T*_*max*_ from 32.6°C (Multi-species estimated) to 36°C (*An. stephensi* estimated) and a decrease in *T*_*min*_ from 19°C (Multi-species estimated) to 15.6°C (*An. stephensi* estimated). The *An. stephensi* estimated model also had a predicted higher *T*_*opt*_ than the previous Multi-species estimated model (*T*_*opt*_; 25.4°C) by 2.2°C. (**Figure 2d**, **SI_Table 6**). Finally, we found differences in the *T*_*opt*_ and *T*_*max*_ of the estimated thermal relationship for the proportion mosquitoes surviving the latency period (exp[*−u*(T)/PDR(T)*]) between the Multi-species estimated and *An. stephensi* estimated models (**SI_Figure 4, SI_Table 7**).

### Seasonal transmission based on temperature suitability varies geographically across relative *R_0_* models

To visualize the differences in model predictions, we created maps illustrating geographic variation in seasonal suitability for *P. falciparum* transmission of the three relative *R*_*0*_*(T)* models along with spatial descriptors of the derived maps (**Figure 3a-c**, **SI_Table 8**). Comparisons are drawn to the Multi-species estimated model to illustrate how environmental suitability predictions may vary across disease systems (**Figure 3d**). The mapped overlay of year-round (12-months) environmental suitability for malaria transmission for all three models highlights the broader geographic extent of temperature suitability in our *An. stephensi-P. falciparum* models, most notably extending northward into India and on the Arabian Peninsula as compared to the previous Multi-species estimated *R*_*0*_(*T*) model (**Figure 3d**, **SI_Table 8**). For example, the Multi-species estimated model predicts India to contain 710,046 km^2^ of endemic area, whereas our two *An. stephensi* models are predicted to contain approximately 3.5 times more endemically suitable area (**SI_Table 8**). Further, Qatar was predicted to be unsuitable for year-round malaria transmission in the Multi-species estimated model but contained a modest area of endemic transmission suitability (11,210 km^2^) with our two *An. stephensi* models. In contrast, the predicted endemically suitable area in Sri Lanka remained largely unaltered among models predictions. In addition, there was a very subtle increase in suitability season in the northern-most regions with the *An. stephensi* lifetime model compared to the *An. stephensi* estimated model (**Figure 3**).

## Discussion

This comprehensive study characterized how mosquito life history traits of an urban Indian malaria vector, *An. stephensi*, were jointly modified by temperature and age to affect the temperature suitability for malaria transmission. This study also tailored current temperature-dependent relative *R*_*0*_ models to *An. stephensi* to evaluate how predictions of environmental suitability for malaria transmission are influenced by: 1) the use of direct observations for mosquito life history traits (e.g. lifespan, lifetime egg production, and biting rate) instead of common proxies currently used in the literature to estimate these traits and 2) the inclusion of *An. stephensi*-specific trait data. We found that in addition to temperature, mosquito age altered the daily proportion of females imbibing a bloodmeal, daily egg production, and daily probability of survival. These results suggest that estimates of these life history traits characterized during a finite portion of a mosquito’s lifespan may be imprecise (24, 31, 32). Importantly, we found large quantitative differences in observed lifetime trait values relative to estimates that suggest absolute transmission potential could differ with mosquito age. A failure to include the effects of mosquito age structure, for example, could have important implications for modeling approaches that predict malaria transmission dynamics. Finally, we determined that the inclusion of *An. stephensi* – *P. falciparum* specific data resulted in qualitatively different temperature-transmission relationships compared to a previous relative *R*_*0*_*(T)* model that used thermal responses from *An. gambiae* and other *Anopheles* and *Aedes* species, ultimately affecting predictions of regional suitability for malaria transmission (31, 32).

Research across a diversity of ectothermic organisms demonstrates that age and temperature both affect development, survival, and reproduction (50–53). In mosquitoes, limited evidence suggests that mosquitoes experience age-related changes in cuticular hydrocarbons, immune function, flight activity, insecticide resistance, biting rates, and survival (61–72). In this study, we observed a decrease in the proportion of females imbibing a blood meal and daily egg production in older *An. stephensi*. For both the proportion of females imbibing blood and survival, the rate of decline occurred faster at increasingly warmer temperatures. Previous work with *An. gambiae* showed an increase in the daily biting rate with age, which is in contrast to our findings (55). It remains unclear whether this is outcome is due to differences in senescence, allocation of resources into different life history tasks across *Anopheles* species, or nutritional conditions. Finally, the daily egg production for *An. stephensi* displayed a unimodal relationship with age, where daily egg production increased to a peak before declining. The time to reach peak values and the subsequent decline occurred earlier in mosquitoes housed at warmer temperatures.

The temperature-sensitive age-dependent mortality rates for mosquito populations are concordant with previous work in laboratory and limited field studies (51, 73, 74). While there is some evidence that long-lived *An. gambiae* cohorts can occur in the field, it is generally assumed that mosquitoes have shorter lifespans in the field than typically observed in controlled laboratory settings (75, 76), and the same may be true for *An. stephensi*. Thus, whether mosquitoes experience senescence in the field remains an open and critical question, primarily due to the logistical difficulties of accurately aging mosquitoes, conducting mark-recapture studies, and controlling for temperature in variable environments (51).

Using direct measurements of an individual’s biting rate, lifetime fecundity, and lifespan instead of common approaches to estimate these traits from truncated portions of a mosquito’s life (e.g., first gonotrophic cycle only) yielded quantitatively, and in some cases qualitatively, different temperature-trait relationships. Our results suggest that previous approaches used to estimate these life history traits in the literature underestimate values for these traits across most temperatures. This could have important ramifications for predicting mosquito population dynamics as well as the effectiveness of mosquito control interventions in the field where thermal conditions vary. Further, imprecise estimates of lifespan can have a compounding effect on predictions of population dynamics and vector-borne disease transmission as it impacts estimates of total reproductive output and the amount of time a mosquito survives past becoming infectious (51, 52, 77). More effort is needed in measuring both lifespan and age-associated changes in life history traits under field settings for important mosquito disease vectors.

By using a temperature-dependent *R*_*0*_ model framework we were able to explore how model parameterization of trait data (estimated vs. observed) influenced the temperature suitability for *P. falciparum* transmission. While there were substantial quantitative differences between directly measured versus estimated lifetime trait values along with qualitative differences in shape of the temperature-dependent functions for biting rate and lifetime egg production, we observed only very subtle differences in the predicted effects of temperature on the environmental suitability of malaria transmission between the *An. stephensi* estimated and *An. stephensi* lifetime models (**Figure 2d, Figure 3**). With the relative *R*_*0*_ approach, absolute differences in predicted temperature-trait and temperature-transmission relationships are masked. One potential ramification of ignoring absolute differences across temperature-trait relationships is that we cannot account for variation in the intensity of malaria transmission with temperature among modeling approaches. For example, even though values of mosquito lifespan differed by up to 2-fold, the predicted temperature-relative *R*_*0*_ relationship was relatively unaffected by whether the temperature-lifespan relationship was parameterized from estimated or directly observed values(**Figure 2**). These results suggest that predictions of seasonal prevalence could be improved in a modelling framework that incorporates the age-structure of mosquito populations.

Additionally, while our *An. stephensi* estimated model was sensitive to lifespan, our *An. stephensi* lifetime model was less so (**SI_Results**, **SI_Figure 2,3**). Thus, the small shifts in the predicted thermal minimum and optimum for relative *R*_*0*_ to cooler temperatures in our *An. stephensi* lifetime model relative to the *An. stephensi* estimated model is primarily driven by the qualitative differences in the temperature-trait relationship between directly observed and estimated biting rate and the proportion of mosquitoes surviving the latency period (**SI_Results**, **Figure 2**, **SI_Figures 2-4**). Differences in the temperature-trait relationship for the proportion of mosquitoes surviving the latency period likely arise between models as the *An. stephensi* lifetime model accounts for mortality rates that vary with age, whereas the *An. stephensi* estimated model assumes a constant mortality rate.

Using *An. stephensi* data dramatically changed the predicted relationship between the environmental suitability of malaria transmission and temperature relative to the previously published Multi-species estimated model based largely on *An. gambiae* and *P. falciparum* (31), suggesting that the thermal limits and optima of relative *R*_*0*_*(T)* models varies across disease systems (26, 78). Specifically, we demonstrate a 3.4°C decrease in the predicted thermal minimum and 3.4°C increase in the thermal maximum for our *An. stephensi* estimated model, as compared to the Multi-species estimated model that used trait responses from multiple *Anopheles* and an *Aedes* species (**Figure 2d**, **SI_Table 4, SI_Table 6**) (32). The increase in environmental suitability at warmer temperatures could be due to differences in physiological constraints of the mosquito vectors investigated. *An. stephensi* may be selected for higher temperature tolerance, as it is found in urban areas in Asia. Thus, due to its geographical location and the urban ‘heat-island effect,’ this species inhabits warmer areas on average than that of the more rural *An. gambiae* (60). Further, differences in *Plasmodium* species and the method of calculating EIP could drive differences between models (79). However, this would not explain the increased suitability at cooler temperatures, which instead suggests a vector or parasite with a higher plasticity in temperature tolerance. Finally, incorporating life history data for *An. stephensi* and *P. falciparum* reduced the credible intervals around the predicted temperature-relative *R*_*0*_ relationship relative to the Multi-species estimated model (**Figure 2d**) (31). In order to further refine temperature suitability predictions for the effective use in vector control and to optimally inform public health strategies there is a strong need for additional research on temperature effects on the basic biology of disease vectors such as mosquitoes.

Accurately predicting malaria transmission depends on variation in other abiotic and biotic factors, as well as socioeconomic factors that determine human exposure to infectious mosquitoes that the *R*_*0*_ approach does not capture. Further, the *R*_*0*_ framework employed in this work is static and does not incorporate the effect of temporal variation in daily or seasonal temperatures, as well as fluctuations in vector and host densities or disease states (e.g., susceptible, exposed, infectious, recovered). However, here we demonstrate that the predicted thermal suitability for endemic malaria transmission in Southeast Asia is more substantial than previously predicted. Additional study limitations associated with the experimental design are presented in **SI_Discussion**. Surprisingly, the predicted temperature-relative *R*_*0*_ relationship and overall land area of endemic environmental suitability were only subtly affected by using common approaches to estimate mosquito lifetime traits (e.g., lifespan, lifetime egg production, and biting rate) versus directly measuring them. However, differences in the overall magnitude of these traits—as opposed to the shapes of their thermal responses—could affect transmission in ways not captured using the relative *R*_*0*_*(T)* approach. This work highlights the importance of integrating data specific to a disease system of interest and underscores the need for more basic research in the field to improve the accuracy of mechanistic transmission models.

## Acknowledgements

The authors would like to thank the CURO (Center for Undergraduate Research Opportunities) program and the NSF REU PopBio (Population Biology of Infectious Diseases) program at the University of Georgia for their assistance in conducting this research. We would also like to thank Drs. Leah Johnson and Melinda Brindley for insightful discussion and Dr. Marta Shocket for deriving the derivates used in the sensitivity analyses for both the *An. stephensi* estimated and lifetime models. This work was supported in-part by the NSF GRFP (Graduate Research Fellowship Program) and a NIH R01 award (1R01AI110793-01A1). EAM and SJR were supported by an NSF EEID grant (DEB 1518681). EAM was also supported by a Hellman faculty fellowship, a Stanford Woods Institute for the Environment – Environmental Ventures Program grant, and a Terman Award.

