## Supplemental Information for "Mosquito species and age influence thermal performance of traits relevant to malaria transmission"

**Methods**

***Rearing of Anopheles stephensi***

*An. stephensi* mosquitoes were sourced from a colony housed at Pennsylvania State University which were originally obtained from the Walter Reed Army Institute of Research (Silver Spring, MD, USA). *An. stephensi* were held at standard insectary conditions (27°C ± 0.5°C, 80% ± 5% relative humidity, and a 12L:12D photoperiod) prior to starting the adult life history experiment. We hatched and transferred 110 larvae to plastic trays (6 Qt., 12.4 cm x 34.6 cm x 21.0 cm) containing 500 mL of distilled water and maintained on a daily regimen of 100 mg ground TetraMin fish flakes. To ensure age-matched individuals were used in the life table experiment, only pupae present on day 9 post-hatch (peak pupal stage) were collected and placed into adult cages. Any pupae remaining after 24 hr were removed. We provided adult mosquitoes with a solution of 5% dextrose and 0.05% para-amino benzoic acid (PABA) upon emergence. For colony maintenance, *An. stephensi* were fed whole human blood (O+, healthy male < 30 years, Interstate Blood Bank, TN, USA).

***Fitting thermal performance curves with a Bayesian approach***

To predict the thermal limits (*T_min_*, *T_max_*) and optimum (*T_opt_*) for each parameter, we used Bayesian inference to fit either a symmetric (quadratic; -*c*(*T-T_min_*)(*T-T_max_*)) or an asymmetric (Briere; *cT*(*T-T_min_*)(*T_max_*-*T*)^1/2^) unimodal non-linear function to each trait versus temperature as in Johnson et al. 2015 (1). We first fit curves with uninformative priors restricted to biologically informed ranges (*T_0_* ~ uniform (0, 24), *T_m_* ~ uniform (25, 45), *c* ~ uniform (0, 1)) , followed by informative priors derived from traits estimated in a previous study (1) (assuming a gamma distribution over each component trait). A direct comparison of each temperature-trait response using either uninformative or informative priors is provided to illustrate the influence of informative priors on our trait fits presented in the main text (**SI_Figure 1**). Both quadratic and Briere functions are truncated at 0 to exclude negative values, where *c* is a fit parameter that controls the shape of each respective function. Models were generated in R using JAGS (2) and the R package <rjags> (3), by running three Markov Chain Monte Carlo simulations for a 5,000-step burn-in followed by 20,000 additional steps, then thinning the posterior samples by saving every eighth sample.

We also defined temperature-trait responses for mosquito and parasite traits not directly measured in this study to assess the impact incorporating multiple trait thermal responses from a single mosquito species (*An. stephensi*), rather than aggregated from several different mosquito species, has on *R_0_*(*T*). *An. stephensi* data from (4) and (5) were used to construct temperature-trait relationships for mosquito development rate (*MDR*), probability of egg to adult survival (*pEA*), *P. falciparum* development rate (*PDR*) and vector competence (*bc*). In contrast, for the Multi-species estimated model we used the thermal relationships defined in (1).

***Derivation of* R_0_(T) *models***

The Ross-Macdonald expression is commonly used to represent *R_0_,* the basic reproductive number, or the number of secondary cases expected to arise from a primary infection given a fully susceptible population (**SI_Equation 1**) (6).

$R_{0}=\sqrt{\frac{Ma^{2}bce^{-\mu EIP}}{Nr\mu}}$	(**SI_Equation 1**)

*R_0_* is composed of parameters for vector abundance, *M*, human host abundance, *N*, the vector biting rate, *a*, vector competence, *bc*, or the product of the proportion of vectors that become infected (*b*) and the proportion of human hosts that become infected (*c*), the vector daily mortality rate, *µ*, human recovery rate, *r*, and the extrinsic incubation period of the pathogen (*EIP*). In accordance with previous studies, we incorporate how these traits are affected by temperature. First, as previously (7), the effect of temperature (*T*) and rainfall (*R*) on mosquito density (*M*) was accounted for by including temperature-sensitivity to mosquito life history traits such as per capita birth rate (*λ*), per capita death rate (*µ*), daily survival probability of larval (*p_L_*), larval development time (*τ_L_*) and the effect of rainfall on as per capita birth rate (*λ*), daily survival probabilities of eggs (*p_E_*), larvae (*p_L_*), and pupae (*p_P_*) as shown in **SI_Equation 2**.

$M\left( R,T \right)=\frac{\lambda\left( R,T \right)}{\mu\left( T \right)} ; \lambda\left( R,T \right)=Bp_{E}(R)p_{L}(R)p_{L}(T)p_{p}(R)/(\tau_{E}+ \tau_{L}\left( T \right)+\tau_{P})$ (**SI_Equation 2**)

In the above expression (**SI_Equation 2**), Parham and Michael 2010 (7) defined *B* to be the number of eggs laid per adult per oviposition, *p_E_*, *p_L_*, and *p_P_* as the daily survival probabilities of eggs, larvae and pupae and *τ_E_*, *τ_L_*, and *τ_P_* as the durations of each of these stages. They assumed *B* to be independent of environmental conditions, development times in each stage to be dependent on temperature only if there was sufficient rainfall to sustain development and independent effects of temperature and rainfall on the daily survival probability of larvae.

Later in (8) and subsequent work (1, 9-12), the dependency of mosquito density (*M*) on rainfall was dropped, all time periods were expressed as daily rates, and fecundity in (7) was interpreted to represent total egg production of an individual female adult (*B*) as opposed to the number of eggs laid per adult per oviposition. Further, as data characterizing daily survival probabilities and development times across temperature for each immature stage are scarce the products of the daily survival probability and the sum of the development times for each stage were replaced with composite traits (*p_EA_* and *τ_EA_*) representing the entire process. Thus, *EIP* is replaced with the reciprocal of the pathogen development rate (*PDR*), and *τ_EA_* is replaced with the reciprocal of the mosquito development rate (*MDR*). Further, as data for *B* were unavailable, *B* was substituted with the expression; *B* = *EFD*/*µ*, where *EFD* is the daily egg production per female and was made to be temperature-dependent (**SI_Equation 3**, **SI_Equation 4**).

$M\left( T \right)=\frac{\lambda\left( T \right)}{\mu\left( T \right)} ; \lambda\left( T \right)=\frac{B{(T)p}_{EA}\left( T \right)}{\tau_{EA}\left( T \right)}=B{(T)p}_{EA}\left( T \right)MDR\left( T \right)= \frac{EFD(T)p_{EA}\left( T \right)MDR\left( T \right)}{\mu\left( T \right)}$

(**SI_Equation 3**)

**SI_Equation 4** is the same formulation used for *R_0_(T)*_estimated_ (**Eq. 1**) in the main text. In **Eq. 1** *a*, *µ*, and *EFD* are marked with an * to denote that the data used to parameterize these traits are estimated (as is commonly done in these models) and do not represent the definitions stated above. For example, biting rate, *a*, is often approximated by using the inverse of the time to the first oviposition instead of directly measuring the number of bites an individual takes in a defined time period.

$M=\frac{{EFD}^{*}(T)pEA(T)MDR(T)}{{\mu^{*}(T)}^{2}} ; R_{0}=\sqrt{\frac{M{a^{*}\left( T \right)}^{2}bc(T)e^{-\mu/PDR(T)}}{Nr\mu^{*}(T)}}$

Simplified	$R_{0}=\sqrt{\frac{{{EFD}^{*}(T)pEA(T)MDR(T)a^{*}\left( T \right)}^{2}bc(T)e^{-\mu/PDR(T)}}{Nr{\mu^{*}(T)}^{3}}}$	(**SI_Equation 4**)

To derive the expression for *R_0_(T)*_lifetime_ (**Eq. 2**) in the main text, we first represented all *µ* terms with lifespan (*lf*; 1/ *µ*). Next, as directly measured *lf*, biting rate (*a*), and the total egg production of an individual female (*B*), the * from these parameters was removed in the expression, and *B* was back substituted in place of *EFD*/*µ.* Finally, to account for the effects of age-variable mortality rates in the proportion of mosquitoes surviving the latency period, *ϒ* is substituted for exp[-*µ*(*T*)*/PDR*(*T*)]. The temperature-trait relationship for the proportion of mosquitoes surviving the latency period, *ϒ*(*T)*, is calculated from the Bayesian fit of the proportion of mosquitoes alive (taken from the Gompertz fits to survivorship from each experimental replicate) upon completion of the predicted extrinsic incubation period (*PDR_50_*(*T*)^-1^ or the amount of days to reach 50% of maximum infectiousness in a mosquito population) of *P. falciparum* at each temperature (5).

${R_{0}(T)}_{lifetime}=\sqrt{\frac{{{a\left( T \right)}^{2}bc\left( T \right)ϒ(T)B(T)pEA(T)MDR(T){lf(T)}^{2}}}{Nr}}$	(**Eq. 2**)

As we do not include values for host-specific traits such as *N* or *r,* and assume these traits are temperature-independent, our static *R_0_* expression is a relative metric of temperature suitability for transmission as opposed to the traditional interpretation of *R_0_* as a metric of disease invasion into a fully susceptible population. Further, absolute values of *R_0_* additionally depend on location-specific factors such as breeding habitat availability, vector biting preference, host availability, disease control efforts, intra- and inter- species interactions, along with additional abiotic factors. Thus, the relative *R_0_* framework is adopted, and the relationship between relative *R_0_* and temperature is used to evaluate the impact of model parameterization on the environmental suitability for *P. falciparum* transmission by *An. stephensi* mosquitoes.

***Sensitivity and uncertainty analyses on* An. stephensi *R_0_(T) models***

To determine if *R_0_(T)* formulation (**Eq. 1** versus **Eq. 2**) affected the sensitivity and uncertainty of *R_0_* to trait parameters, we performed two types of sensitivity analyses and an uncertainty analysis on our two *An. stephensi* *R_0_(T)* models. For a similar analysis on the Multi-species estimated model see (1). Specifically, our *An. stephensi* lifetime model contains one less *µ* term due to the substitution of lifetime egg production (*B*) for *EFD*/*µ* and allows age-dependent daily mortality in the proportion of mosquitoes surviving the latency period (*ϒ*). First, to illustrate the degree to which a small change in trait *x* affects *R_0_* at a given temperature (*T*), a model derivative (d*R_0_*/d*x*) was divided by *R_0_* for each trait *x*, to give d*R_0_*/ (*R_0_* d*x*) or the standardized sensitivity of *R_0_* to trait *x* across all temperatures. Second, to demonstrate the impact of temperature sensitivity of a given trait *x* on *R_0_*, relative *R_0_(T)* was calculated with each trait held at a constant value and allowing the other parameters to vary with temperature. Finally, we estimated the uncertainty in *R_0_* introduced through uncertainty in each temperature-trait relationship. To do this we calculated *R_0_(T)* by allowing each trait *x* to assume its full posterior distribution *x(T)* while setting all other traits to their posterior median thermal responses. Then, we calculated the width of the 95% credible interval on *R_0_* at each temperature to estimate the partial uncertainty with respect to *x*. To determine the full uncertainty in relative *R_0_*(*T*)*,* the width of the 95% credible interval of *R_0_* at each temperature was calculated by allowing all parameters to assume their full posterior distribution. We then divided partial uncertainty for each trait by full uncertainty to estimate the proportion of total uncertainty in relative *R_0_* that is driven by each trait *x* at each temperature *T*.

**Results**

***Model formulation affects the sensitivity and uncertainty of relative* R_0_(T) *to trait parameters***

We conducted two types of sensitivity and an uncertainty analysis to determine if the differences in *R_0_(T)* formulations between the *An. stephensi* lifetime and *An. stephensi* estimated models affected the relative sensitivity and uncertainty of *R_0_* to different trait parameters. Differences in model formulation did alter the thermal sensitivity of *R_0_* to each trait parameter as well as which traits contributed most to uncertainty in *R_0_* across the thermal spectrum (**SI_Figure 2, SI_Figure 3**). However, direction comparison of the relative sensitivity and uncertainty of a given trait between *R_0_* models should be interpreted cautiously, as traits are not equally represented between models. *R_0_(T)* was sensitive to lifespan (*lf*) and biting rate (*a*) in both *An. stephensi* models; however, the *An. stephensi* lifetime model exhibited less sensitivity to lifespan (*lf*) than the *An. stephensi* estimated model (*An. stephensi* lifetime; **SI_Figure 2B,C** & *An. stephensi* estimated; **SI_Figure 2B,C**). Much of our *An. stephensi* lifetime model uncertainty was attributed to vector competence (*bc*; across all temperatures), followed by lifetime egg production (*B*, intermediate to warm temperatures), and lifespan (*lf*, warm temperatures) (**SI_Figure 2D**). In contrast, uncertainty around the temperature-trait relationship that contributed the largest proportion of uncertainty in the *An. stephensi* estimated model was from estimated (*lf**) at temperatures greater than 20°C (**SI_Figure 3D**). Vector competence (*bc*, across all temperatures), estimated biting rate (*a**, at intermediate temperatures), parasite development rate (*PDR*, temperatures below 20^o^C), and vector competence (bc, across all temperatures) also contributed uncertainty in our *An. stephensi* estimated model (**SI_Figure 3D)**.

| **Tables**  **Supplemental Table 1: Generalized linear mixed model selection** | | | | | | | | | |  |
| --- | --- | --- | --- | --- | --- | --- | --- | --- | --- | --- |
| ***Daily Bite Rate*** | |  |  |  | |  |  |  |  |  |
| **Model Name** | **Formula** | **(Intercept)** | **Dscale** | | **Tscale** | **Dscale:Tscale** | **Class** | **df** | **LogLik** | **AICc** |
| ***Feed.m16*** | ***Feed ~Tscale * Dscale + (1\|Female)*** | ***-0.622337*** | ***NA*** | | ***0.4576745*** | ***NA*** | ***glmerMod*** | ***3*** | ***-6952.4*** | ***13910.82*** |
| Feed.m20 | Feed ~ Tscale + (1\|Donor) + (1\|Block) | -0.6262926 | NA | | 0.464569 | NA | glmerMod | 4 | -6951.7 | 13911.37 |
| Feed.m21 | Feed ~ Tscale + (1\|Donor) + (1\|Female) | -0.6223341 | NA | | 0.4576742 | NA | glmerMod | 4 | -6952.4 | 13912.82 |
| Feed.m11 | Feed ~ Tscale * Dscale + (1\|Donor) | -0.6171723 | 0.01413019 | | 0.4702643 | 0.030024438 | glmerMod | 5 | -6951.6 | 13913.11 |
| Feed.m6 | Feed ~ Tscale * Dscale + (1\|Block) + (1\|Donor) | -0.6205138 | 0.00859818 | | 0.4709623 | 0.022081056 | glmerMod | 6 | -6951.3 | 13914.63 |
| Feed.m7 | Feed ~ Tscale * Dscale + (1\|Donor) + (1\|Female) | -0.6171683 | 0.01412949 | | 0.4702615 | 0.030025017 | glmerMod | 6 | -6951.6 | 13915.11 |
| Feed.m1 | Feed ~ Dscale * Tscale + (1\|Block) + (1\|Female) + (1\|Donor) | -0.6205126 | 0.008598 | | 0.4709621 | 0.022082085 | glmerMod | 7 | -6951.3 | 13916.63 |
| Feed.m2 | Feed ~ Tscale + (1\|Block) | -0.6373151 | NA | | 0.4544688 | NA | glmerMod | 3 | -6961 | 13928.03 |
| Feed.m10 | Feed ~ Tscale * Dscale + (1\|Block) | -0.6340627 | 0.04348144 | | 0.4685816 | 0.008365942 | glmerMod | 5 | -6959 | 13928.09 |
| Feed.m4 | Feed ~ Tscale * Dscale + (1\|Block) | -0.6340627 | 0.04348144 | | 0.4685816 | 0.008365942 | glmerMod | 5 | -6959 | 13928.09 |
| Feed.m5 | Feed ~ Tscale * Dscale + (1\|Block) + (1\|Female) | -0.6340693 | 0.04348068 | | 0.4685841 | 0.008369095 | glmerMod | 6 | -6959 | 13930.09 |
| Feed.m9 | Feed ~ Tscale * Dscale + (1\|Block)+ (0+Day\|Female) | -0.6340625 | 0.04348065 | | 0.4685803 | 0.008364724 | glmerMod | 6 | -6959 | 13930.09 |
| Feed.m14 | Feed ~ Tscale + Dscale | -0.6324588 | 0.04406536 | | 0.4422736 | NA | glm | 3 | -6966.7 | 13939.37 |
| Feed.m15 | Feed ~ Tscale * Dscale | -0.6239916 | 0.0485246 | | 0.4538526 | 0.032975864 | glm | 4 | -6965.7 | 13939.46 |
| Feed.m8 | Feed ~ Tscale * Dscale + (1\|Female) | -0.6239916 | 0.0485246 | | 0.4538526 | 0.032975865 | glmerMod | 5 | -6965.7 | 13941.46 |
| Feed.m12 | Feed ~ Tscale | -0.6317109 | NA | | 0.429047 | NA | glm | 2 | -6968.8 | 13941.63 |
| Feed.m18 | Feed ~ Tscale + (1\|Female) | -0.6341786 | NA | | 0.428488 | NA | glmerMod | 3 | -6968.7 | 13943.43 |
| Feed.m19 | Feed ~ Dscale + (1\|Female) | -0.6070619 | -0.0613142 | | NA | NA | glmerMod | 3 | -7109.5 | 14224.95 |
| Feed.m17 | Feed ~Tscale * Dscale + (Dscale\|Female) + (1\|Block) | -0.4687329 | -0.2100982 | | NA | NA | glmerMod | 3 | -7128.4 | 14262.8 |
| Feed.m3 | Feed ~ Dscale + (1\|Block) | -0.4208503 | -0.0511823 | | NA | NA | glmerMod | 3 | -7134.7 | 14275.47 |
| Feed.m13 | Feed ~ Dscale | -0.6075728 | -0.0832071 | | NA | NA | glm | 2 | -7187 | 14378.04 |
| ***Daily Egg Production*** | |  |  | |  |  |  |  |  |  |
| **Model Name** | **Formula** | **(Intercept)** | **Dscale** | | **Tscale** | **Dscale:Tscale** | **Class** | **df** | **LogLik** | **AICc** |
| ***EFD.m4*** | ***EFD.t ~ Tscale * Dscale + (1\|Block)*** | ***2.098319*** | ***0.22191793*** | | ***0.3849332*** | ***-0.0441443*** | ***glmerMod*** | ***6*** | ***-1552.2*** | ***3116.481*** |
| EFD.m5 | EFD.t ~ Tscale * Dscale + (1\|Donor) | 2.098323 | 0.22191997 | | 0.3849355 | -0.04414379 | glmerMod | 6 | -1552.2 | 3116.481 |
| EFD.m1 | EFD.t ~ Dscale * Tscale + (1\|Block) + (1\|Donor) | 2.098319 | 0.22192064 | | 0.384934 | -0.04414629 | glmerMod | 7 | -1552.2 | 3118.527 |
| EFD.m2 | EFD.t ~ Tscale + (1\|Block) | 2.12791 | NA | | 0.308035 | NA | glmerMod | 4 | -1555.9 | 3119.854 |
| EFD.m6 | EFD.t ~ Tscale + (1\|Donor) | 2.127909 | NA | | 0.3080359 | NA | glmerMod | 4 | -1555.9 | 3119.854 |
| EFD.m3 | EFD.t ~ Dscale + (1\|Block) | 2.167493 | 0.05679343 | | NA | NA | glmerMod | 4 | -1564.5 | 3136.996 |
| EFD.m7 | EFD.t ~ Dscale + (1\|Donor) | 2.167492 | 0.05679326 | | NA | NA | glmerMod | 4 | -1564.5 | 3136.996 |

| **Supplemental Table 2: Temperature and day effects on trait values** | | | | | | | |
| --- | --- | --- | --- | --- | --- | --- | --- |
| **Trait** | **Model type** | **Random Effects** | **Family** | **Fixed Effects** | **χ²** | **df** | **p** |
| Proportion of females that imbibed blood | GLMM | Female | Binomial (link = 'logit') | ***Temperature*** | ***421.524*** | ***1*** | ***<0.001*** |
|  |  |  |  | ***Day*** | ***4.2794*** | ***1*** | ***0.03858*** |
|  |  |  |  | Temperature*Day | 1.908 | 1 | 0.16718 |
| Daily egg production | GLMM | Donor | Gamma  (link = 'log') | ***Temperature*** | ***25.3993*** | ***1*** | ***<0.001*** |
|  |  |  |  | ***Day*** | ***7.1834*** | ***1*** | ***0.0074*** |
|  |  |  |  | Temperature*Day | 0.2546 | 1 | 0.6138 |
| Survivorship | Survival | Block | Log-rank test | ***Temperature*** | ***220*** | ***5*** | ***<0.001*** |
| *Distribution* | Exponential | Gamma | ***Gompertz*** | Log-normal | Weibull | | |
| *AIC* | 2979 | 2816 | ***2758*** | 2913 | 2773 | | |

| **Supplemental Table 3: Survival distribution AIC values by temperature treatment** | | | | | |
| --- | --- | --- | --- | --- | --- |
| **Temperature** | **Survival Distribution** | | | | |
|  | Exponential | Gamma | ***Gompertz*** | Log-normal | Weibull |
| 16 | 453 | 437 | ***431*** | 451 | 433 |
| 20 | 451 | 428 | ***418*** | 439 | 422 |
| 24 | 487 | 438 | 438 | 441 | ***436*** |
| 28 | 759 | 720 | ***693*** | 746 | 708 |
| 32 | 447 | 437 | ***426*** | 456 | 432 |
| 36 | 380 | 346 | ***325*** | 360 | 336 |

| **Supplemental Table 4: Source of data included in *R_0_(T)* models** | | | |
| --- | --- | --- | --- |
| **Model** | **Parameter Source** | | |
|  | *a, B, lf* | *bc, PDR* | *MDR, pEA* |
| *An. stephensi^a^* | **this study** | *Shapiro et al. 2017* | *Paaijmans et al. 2013* |
| Multi-vector*^b^* | *Johnson et al. 2015* | *Johnson et al. 2015* | *Johnson et al. 2015* |
| Parameters: daily biting rate (*a*), lifetime egg production (*B*), lifespan (*lf*), vector competence (*bc*), parasite development rate (*PDR*), mosquito development rate (*MDR*), and the probability of egg to adult survival (*pEA*).^a^denotes *R_0_(T)* models describing the environmental suitability of transmission of *P. falciparum* via *An. stephensi*. ^b^indicates the previous *R_0_(T)* model intended to describe the environmental suitability of the transmission of *P. falciparum* via *An. gambiae*. | | | |

| **Supplemental Table 5: Thermal thresholds of temperature-trait relationships of lifetime values** | | | | | | |
| --- | --- | --- | --- | --- | --- | --- |
| **Trait** | **Lifetime Values** | **Function** | **Thermal threshold (95% CI^a^)** | | | |
|  |  |  | *T_min_* (°C) | *T_opt_ (*°C) | *T_max_* (°C) | *T_breadth_* (°C)^b^ |
| biting rate | observed (*a*) | Briere | 2.6 (0.4-5) | 33.6 (32.8-35) | 41.8 (40.4-43.4) | 39.2 (35.4-43) |
|  | estimated (*a**) |  | 10.2 (6.8-13) | 37.8 (34.8-41) | 44.2 (42.4-45) | 34 (29.4-38.2) |
| lifespan | observed (*lf*) | Quadratic | 1.2 (0-3.6) | 19.6 (18.8-20.6) | 37.8 (37-38.6) | 36.6 (33.4-38.6) |
|  | estimated (*lf**) |  | 1 (0-3.8) | 19.6 (18.4-20.8) | 37.8 (36.4-39.2) | 36.8 (32.6-39.2) |
| lifetime egg production | observed (*B*) | Quadratic | 12.2 (10.6-13.6) | 25.4 (24.4-26.2) | 38.4 (36.8-39.8) | 26.2 (23.2-29.2) |
|  | estimated (*B**) |  | 13.2 (11.2-15) | 27.6 (26.8-28.4) | 39 (37.6-40.6) | 25.8 (22.5-29.4) |
| Thermal threshold values are based on median model outputs. ^a^CI represents the credible interval of Bayesian fits. ^b^*T_breadth_* is the range of temperatures that the trait-function is greater than 0 (*T_max_*-*T_min_*). Data source is from the lifetable experiment conducted in this study. | | | | | | |

| **Supplemental Table 6: Thermal thresholds of relative *R_0_(T)* models** | | | | |
| --- | --- | --- | --- | --- |
| ***R_0_(T)* model** | **Thermal threshold (95% CI)** | | | |
|  | *T_min_* (°C) | *T_opt_ (*°C) | *T_max_* (°C) | *T_breadth_* (°C)^b^ |
| *An. stephensi* lifetime | 15.2 (14.2-16.4) | 26.6 (26.4-27) | 36 (35.4-36.2) | 20.8 (19-22) |
| *An. stephensi* estimated | 15.6 (14.2-16.4) | 27.6 (27-28) | 36 (35.4-36.2) | 20.4 (19-22) |
| Multi-species estimated | 19 (15.8 - 23) | 25.4 (24-27) | 32.6 (29.8-34.6) | 13.6 (6.8-18.8) |
| Thermal threshold values are based on scaled and rounded median model outputs. ^a^CI represents the credible interval. ^b^*T_breadth_* is the range of temperatures that *R_0_(T)* > 0 (*T_max_*-*T_min_*). | | | | |

| **Supplemental Table 7: Thermal thresholds of *Ƴ(T)* across *R_0_(T)* models** | | | | |
| --- | --- | --- | --- | --- |
| **Model** | **Thermal threshold (95% CI)** | | | |
|  | *T_min_* (°C) | *T_opt_ (*°C) | *T_max_* (°C) | *T_breadth_* (°C)^b^ |
| *An. stephensi* lifetime | 8.2 (5.2-11) | 25.8 (24.4-27) | 43.6 (41.6-45) | 35.4 (30.6-39.8) |
| *An. stephensi* estimated | 14.2 (11.8-16.2) | 27.6 (26.6-28.2) | 37 (35.4-38) | 22.8 (19.2-26.2) |
| Multi-species estimated | 14 (13-15) | 24.8 (23.6-26) | 34.4 (33.6-34.8) | 20.4 (18.6-21.8) |
| Thermal threshold values are based on scaled median model outputs. ^a^CI represents the credible interval. ^b^*T_breadth_* is the range of temperatures that *Ƴ(T)* > 0 (*T_max_*-*T_min_*). | | | | |

| **Supplemental Table 8: Endemic area** | | | |
| --- | --- | --- | --- |
| **Country** | **Endemic Area (km^2^)** | | |
|  | *An. stephensi* lifetime | *An. stephensi* estimated | Multi-species estimated |
| India | 2,428,528 | 2,372,906 | 710,046 |
| Oman | 299,702 | 300,880 | 103,645 |
| Pakistan | 126,856 | 120,635 | 547 |
| Qatar | 11,210 | 11,210 | 0 |
| Sri Lanka | 65,996 | 65,912 | 64,299 |
| U.A.E. | 79,934 | 79,856 | 78 |

**Figures**

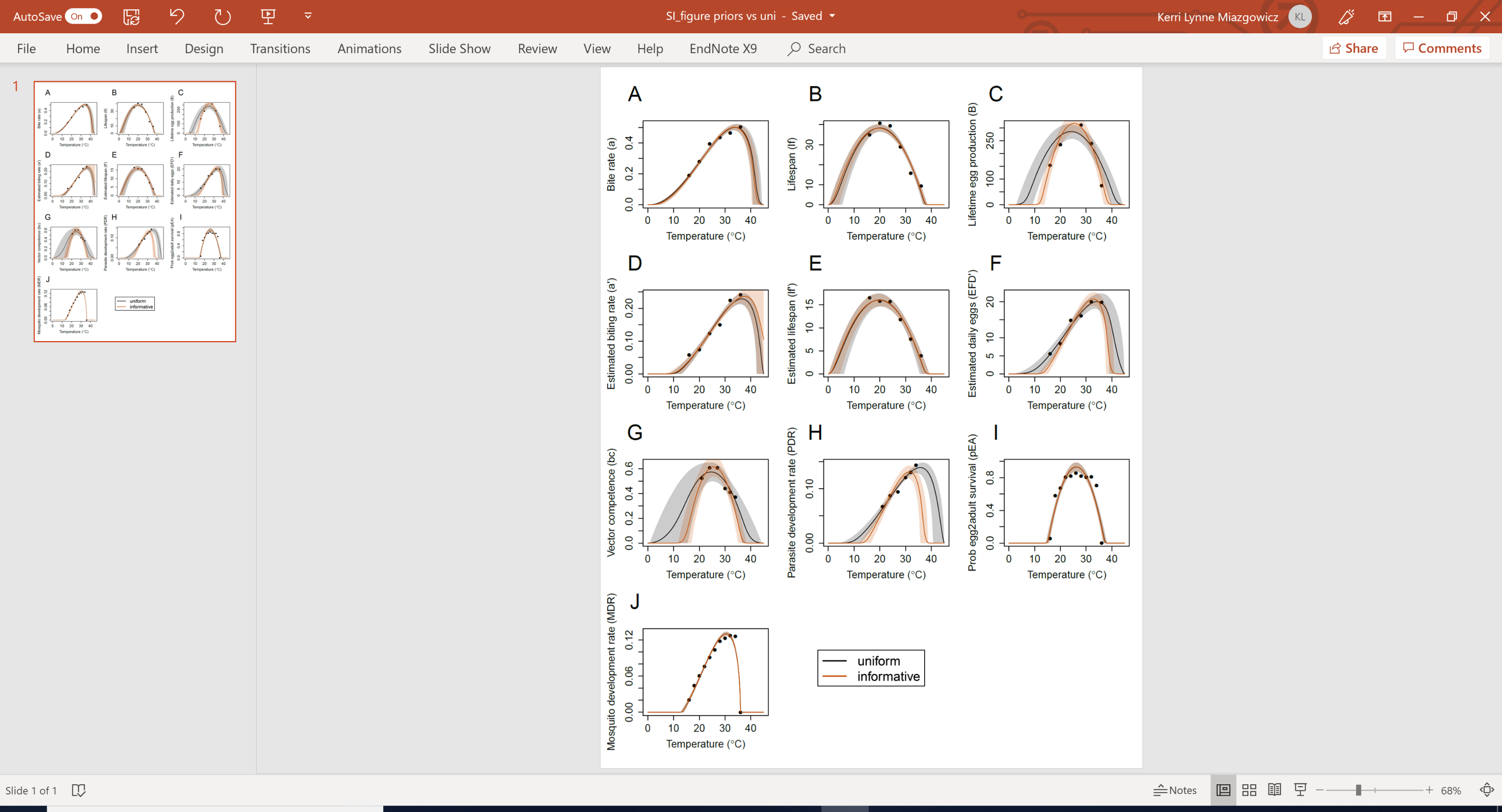
**Supplemental Figure 1.** **Direct Comparison of Uniform to Informative Priors.** Direct comparison on how the use of uniform (black) or informative (orange) priors influences the temperature-trait relationship and 95% credible intervals for (**A**) bite rate (*a*), (**B**) lifespan (*lf*), (**C**) lifetime egg production (*B*), (**D**) estimated biting rate (*a**), (**E**) estimated lifespan (*lf**), (**F**) estimated daily egg production (*EFD**), (**G**) vector competence (*bc*), (**H**) parasite development rate (*PDR*), (**I**) probability of egg to adult survival (*pEA*), and (**J**) mosquito development rate (*MDR*). Trait thermal performance curves from Johnson et al. 2015 were used as informative priors.

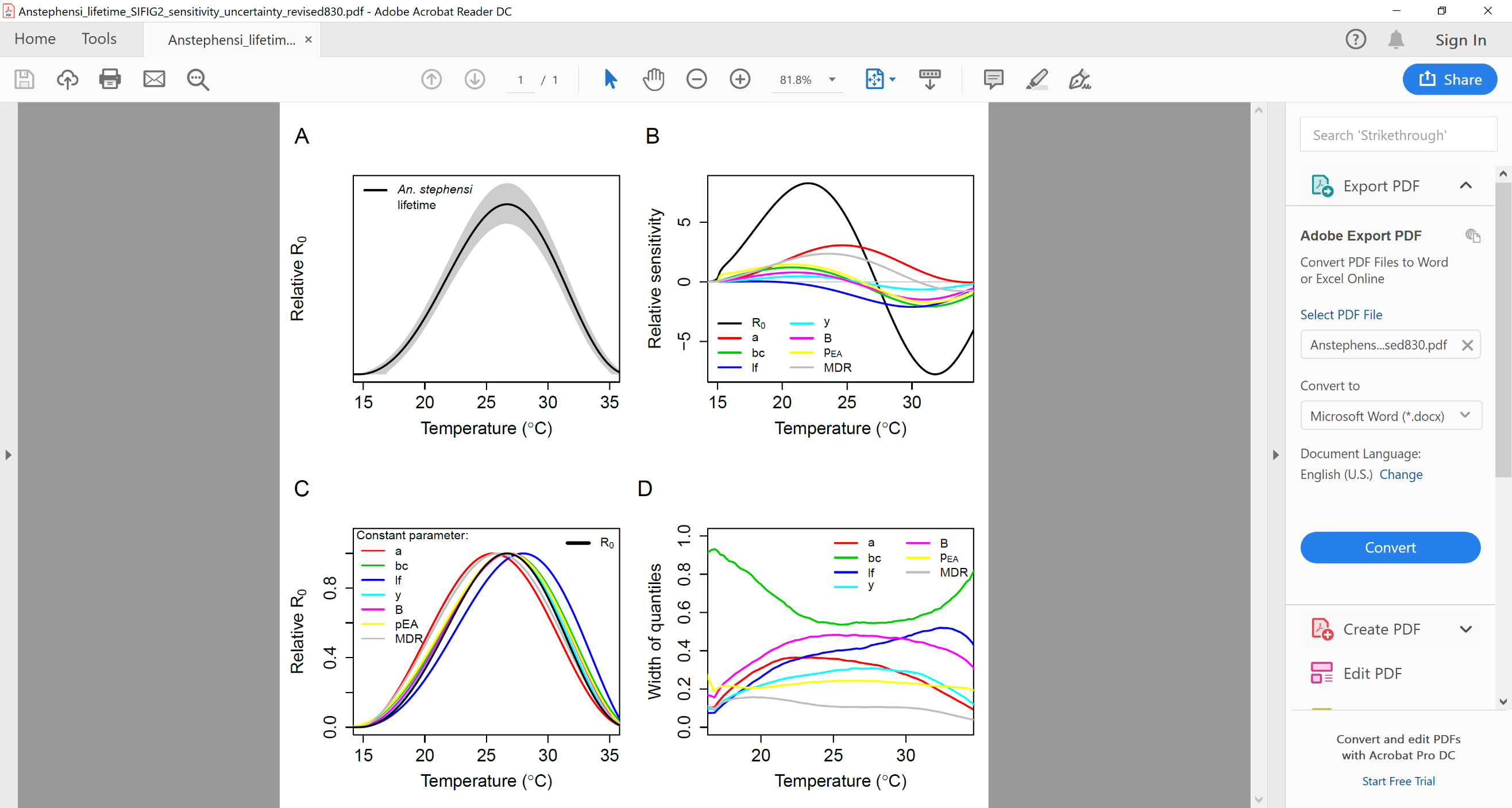
**Supplemental Figure 2. Sensitivity and Uncertainty Analysis on *An. stephensi* lifetime *R_0_(T)* model.** Relative *R_0_(T)* *An. stephensi* lifetime model (**A**) with mean model outputs (solid black line) and 95% credible intervals (dashed black lines). Sensitivity analysis based on formula derivatives for *An. stephensi* lifetime *R_0_(T)* where for each trait x, d*R_0_*/d*x* was divided by *R_0_*, to give d*R_0_*/*R_0_*d*x*, or the standardized sensitivity of *R_0_* to a parameter *x*, across all temperatures (**B**). A second sensitivity analysis based on setting each parameter constant across temperature for *An. stephensi* lifetime where relative *R_0_(T)* was calculated with a single trait held constant and allowing the other parameters to vary with temperature (**C**). Uncertainty analysis on *An. stephensi* lifetime (**D**). For the uncertainty analysis, each trait *x* was allowed to assume its full posterior distribution *x(T)*, while setting all other traits to their posterior median thermal responses and calculating *R_0_*. Partial uncertainty with respect to trait *x* is the width of the 95% credible interval on *R_0_(T)* at each temperature. Full uncertainty was calculated by allowing all parameters to assume their full posterior distribution and calculating the width of the 95% credible interval of *R_0_(T)* at each temperature. Partial uncertainty for each trait divided by full uncertainty of *R_0_(T)* gives the proportion of total uncertainty in *R_0_(T)* that is driven by each trait *x* at each temperature, T.

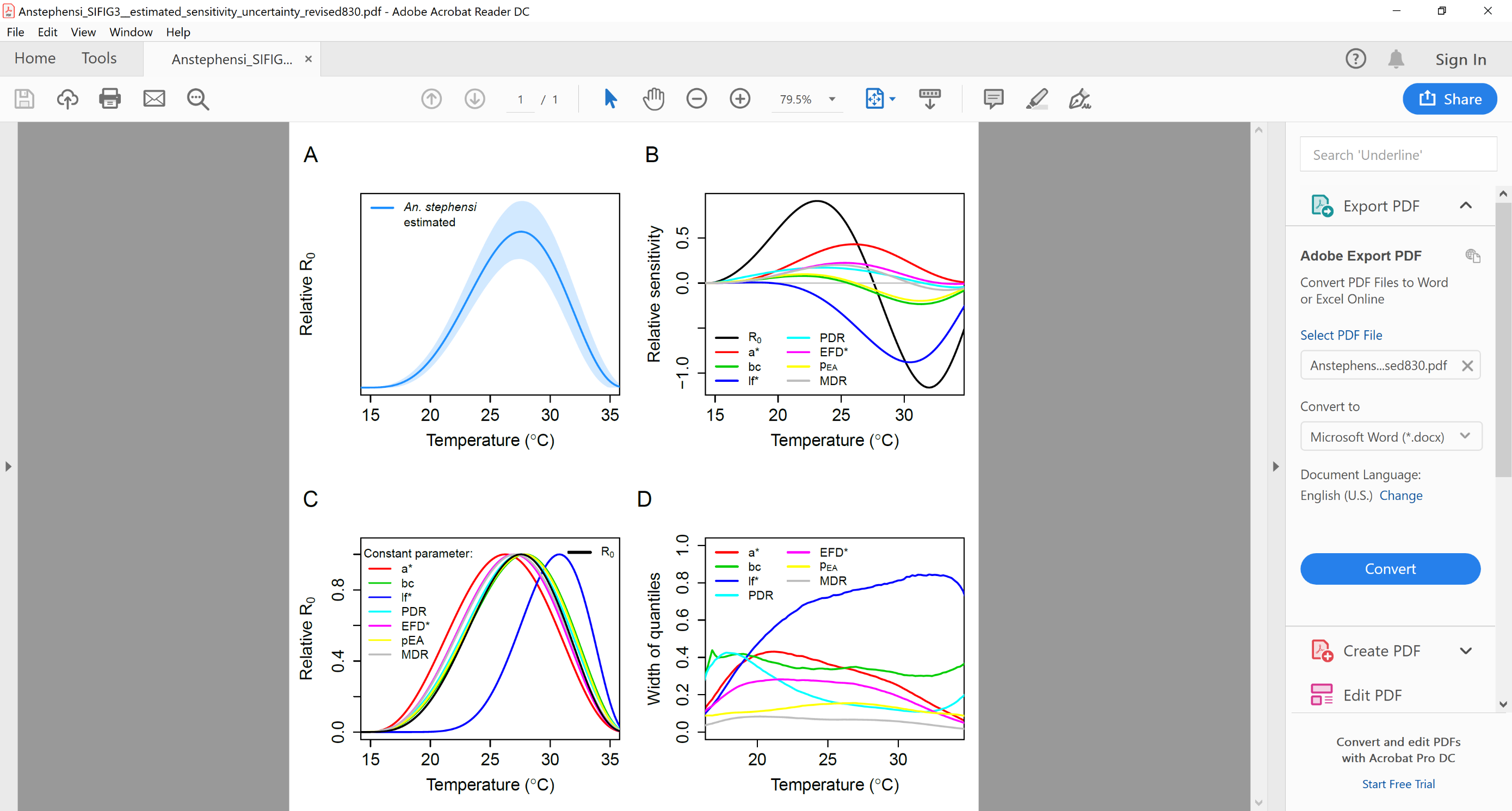
**Supplemental Figure 3. Sensitivity and Uncertainty Analysis on *An. stephensi* estimated *R_0_(T)* model.** Relative *R_0_(T)* *An. stephensi* estimated model (**A**) with mean model outputs (solid black line) and 95% credible intervals (dashed black lines). Sensitivity analysis based on formula derivatives for *An. stephensi* estimated *R_0_(T)* where for each trait x, d*R_0_*/d*x* was divided by *R_0_*, to give d*R_0_*/*R_0_*d*x*, or the standardized sensitivity of *R_0_* to a parameter *x*, across all temperatures (**B**). A second sensitivity analysis based on setting each parameter constant across temperature for *An. stephensi* estimated where relative *R_0_(T)* was calculated with a single trait held constant and allowing the other parameters to vary with temperature (**C**). Uncertainty analysis on *An. stephensi* estimated (**D**). For the uncertainty analysis, each trait *x* was allowed to assume its full posterior distribution *x(T)*, while setting all other traits to their posterior median thermal responses and calculating *R_0_*. Partial uncertainty with respect to trait *x* is the width of the 95% credible interval on *R_0_(T)* at each temperature. Full uncertainty was calculated by allowing all parameters to assume their full posterior distribution and calculating the width of the 95% credible interval of *R_0_(T)* at each temperature. Partial uncertainty for each trait divided by full uncertainty of *R_0_(T)* gives the proportion of total uncertainty in *R_0_(T)* that is driven by each trait *x* at each temperature, T.

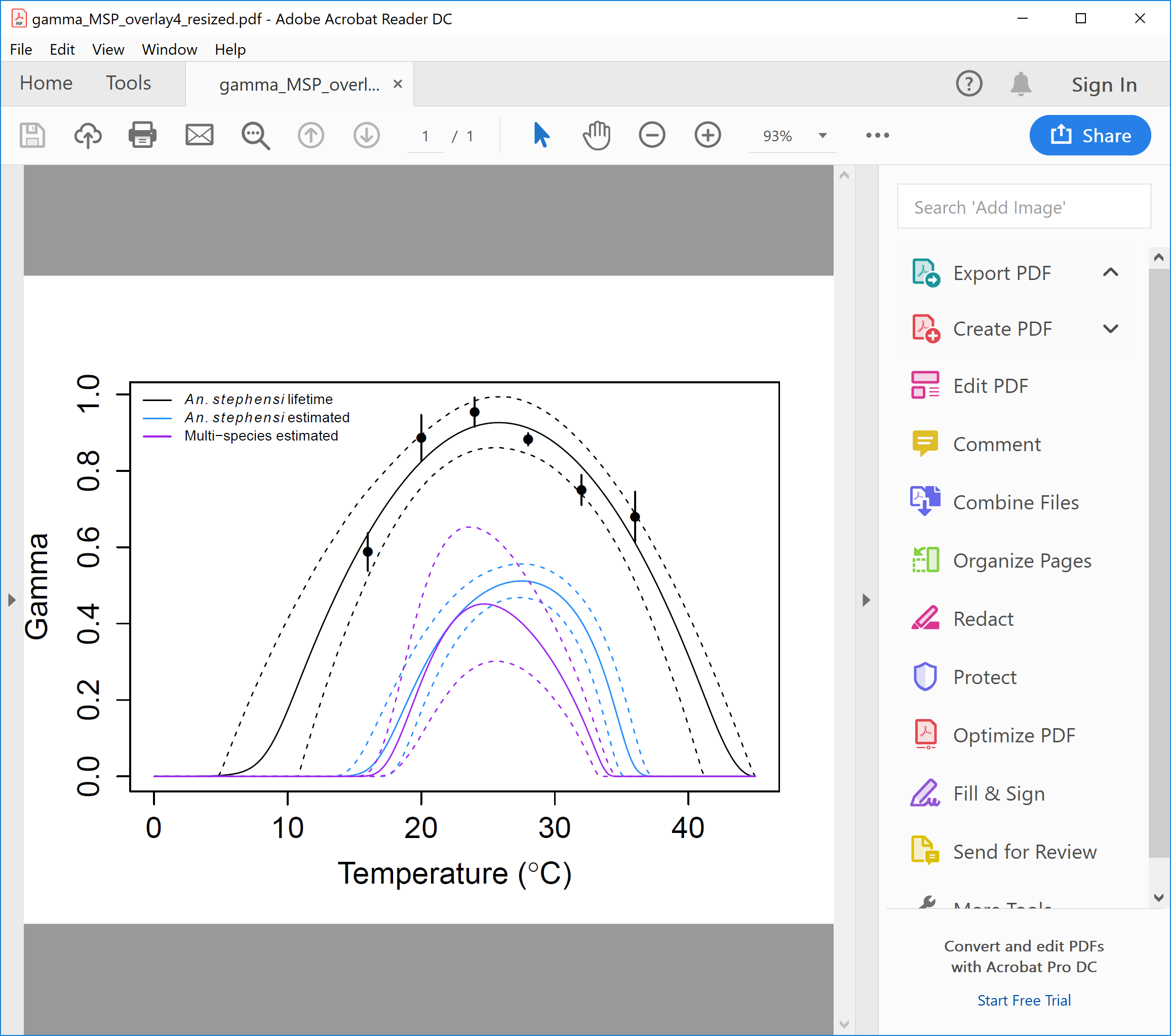
**Supplemental Figure 4. Overlay of gamma (*Ƴ*), the proportion of mosquitoes surviving past the latency period used in the *R_0_(T)* models.** The *An. stephensi* lifetime model incorporates the temperature-trait relationship for the parameter gamma, *ϒ*(*T*), the proportion of mosquitoes surviving the latency period*.* We generated *ϒ(T)* for the *An. stephensi* lifetime model by fitting a quadratic function with Bayesian inference over the proportion of mosquitoes alive (taken from the Gompertz fits to survivorship from each experimental replicate) upon completion of the predicted extrinsic incubation period (*PDR*(*T*)^-1^) of *P. falciparum* at each temperature (as fit from the EIP_50_ values from (5)). *ϒ(T)* for the *An. stephensi* and Multi-species estimated models can be derived indirectly from the following expression; *ϒ*(*T*)= exp[-*µ*(T)/PDR*(*T*)]. In the *An. stephensi* estimated model *µ** was calculated by assuming an exponential function over a truncated portion of the Kaplan-Meier survival estimates as specified in (8). In the Multi-species estimated model *Ƴ(T)* was also indirectly calculated using the expression *ϒ*(*T*)= exp[-*µ*(T)/PDR*(*T*)], however, *µ*(T)* and *PDR(T)* are from the fits generated in Johnson et al. 2015.

**Discussion**

***Study limitations***

Several methodological choices made throughout this study likely influenced the outcome of this experiment. First, we measured mosquito life history traits at constant temperatures. Mosquitoes and their pathogens live in thermally fluctuating environments and life history trait values derived at constant temperatures often differ from those measured under temperature fluctuation (4, 13, 14). However, it is not feasible to directly measure trait performance for all possible permutations of temperature fluctuations a mosquito could encounter, while the characterization of temperature-trait responses at constant temperatures is tractable. Second, all larvae were reared at the same standard temperature (27^o^C) and not at the temperature adult females were eventually held in order to meet the data inclusion criteria outlined in Mordecai et al 2013 (1, 8). These inclusion criteria were adopted to isolate the effects of temperature on adult mosquito traits from carry-over effects of larval rearing temperature. Yet, numerous studies across different mosquito genera have shown the importance of carry-over effects of larval environments on aspects of adult life history (e.g., body size, survival, reproduction, biting rates, and vector competence) (15). Third, adult mosquitoes were not provided sugar during this experiment. Nutrition treatments can affect adult female life history such as biting rate, fecundity, and survival (16-19). However, the extent of sugar feeding of *An. stephensi* in highly urbanized centers is unknown. Finally, these experiments were conducted with a laboratory strain of *An. stephensi* due to the logistical difficulties of acquiring field-based *An. stephensi* from malaria endemic regions. Evidence from other ectotherm and dipteran systems, including mosquitoes, suggests thermal adaptation does occur to local environments (20-24). In addition to the possibility of local adaptation, there is also substantial evidence across different mosquito genera that the ability to acquire and transmit pathogens can also vary across mosquito populations (25-27). Thus, wild populations of *An. stephensi* could have different temperature-trait responses and predicted environmental suitability for *P. falciparum* transmission than characterized in this study (28, 29). However, limited evidence comparing thermal performance in long standing colonies to recently field derived colonies suggests the thermal performance characterized from laboratory colonies might be reasonable reflections of thermal performance in the field for some mosquito species, though more work is needed across a diversity of systems to confirm this (29).
